# The influence of prosocial priming on visual perspective taking and automatic imitation

**DOI:** 10.1101/333880

**Authors:** Rachel Newey, Kami Koldewyn, Richard Ramsey

## Abstract

Imitation and perspective taking are core features of non-verbal social interactions. We imitate one another to signal a desire to affiliate and consider others’ points of view to better understand their perspective. Prior research suggests that a relationship exists between prosocial behaviour and imitation. For example, priming prosocial behaviours has been shown to increase imitative tendencies in automatic imitation tasks. Despite its importance during social interactions, far less is known about how perspective taking might relate to either prosociality or imitation. The current study investigates the relationship between imitation and perspective taking by testing the extent to which these skills are similarly modulated by prosocial priming. Across all experimental groups, a surprising ceiling effect emerged in the perspective taking task (the Director’s Task), which prevented the investigation prosocial priming on perspective taking. A comparison of other studies using the Director’s Task shows wide variability in accuracy scores across studies and is suggestive of low task reliability. In addition, despite using a high-power design, and contrary to three previous studies, no effect of prosocial prime on imitation was observed. Meta-analysing all studies to date suggests that the effects of prosocial primes on imitation are variable and could be small. The current study, therefore, offers caution when using the Director’s Task as a measure of perspective taking with adult populations, as it shows high variability across studies and may suffer from a ceiling effect. In addition, the results question the size and robustness of prosocial priming effects on automatic imitation. More generally, by reporting null results we hope to minimise publication bias and by meta-analysing results as studies emerge and making data freely available, we hope to move towards a more cumulative science of social cognition.

## Introduction

Social interactions involve a number of cognitive processes and behaviours, including imitation and perspective taking. While both of these social skills have been studied extensively in isolation, the relationship between imitation and perspective taking has received less attention. In addition, although social context modulates imitation [1, 2, 3] much less is known regarding how social context influences perspective taking. The current study investigates the relationship between imitation and perspective taking by testing the extent to which these skills are similarly modulated by prosocial priming.

Imitation, or mimicry, is a common occurrence during social encounters, and involves spontaneous copying of others’ actions and gestures [4]. Although such behaviour rarely reaches conscious awareness for either interaction partner, it subconsciously signals a desire to affiliate and build rapport [5]. For example, people who are imitated are bigger tippers [6], donate more to charity [7], engage in prosocial behaviours [7, 8, 9, 10] and indicate liking people who imitate them more than those that do not [6]. Clearly, then, imitation can play an important role in guiding social interactions. To clarify the role imitation can play across different social contexts, recent research has started to identify its antecedents [1, 2]. For example, prosocial priming can increase imitative behaviour [1]. Thus, there exists a bi-directional relationship between imitation and prosociality; those who are imitated behave more prosocially and those who are prosocially primed imitate more. These studies have all employed observational techniques to study imitation, with the measurement being the frequency of observed copying behaviours during live social interaction.

Other researchers devised an index of imitation that is based on reaction time measures. The automatic imitation task [11,12] is an example of a stimulus-response-compatibility (SRC) paradigm, referring to the fact that people cannot help but be affected by the presence of an irrelevant stimulus feature [13,14]. In one well-established automatic imitation task, individuals are instructed to respond to a number cue by lifting their index or middle finger. Concurrently, participants either observe a congruent or incongruent finger movement. Reactions times (RT) are longer in the incongruent compared to congruent condition and this difference is thought to signify the cost of inhibiting an imitative response [1, 15]. Here, then, imitation is captured as the time it takes to suppress the urge to copy an observed action and prioritise one’s own action. The tendency towards imitation (incongruent RT less congruent RT) will hereafter be referred to as the congruency effect.

A handful of studies have explored the effects of prosocial priming on automatic imitation [16, 17, 18]. Priming is thought to operate by subtly triggering a goal that unconsciously guides behaviour [19]. The logic of these studies is that a prosocial prime would activate a goal to affiliate and that this goal would be achieved through increasing the tendency to imitate [16]. Despite using slightly different variants of the automatic imitation task and different experimental designs (see Method section, Table 3), each study reported an effect of prosocial priming on automatic imitation; priming increased the congruency effect. More specifically, the prosocial prime had to be self-related to increase imitation (e.g., ‘I am prosocial’); when using third person primes (e.g., ‘Alex is prosocial’) the congruency effect did not differ from controls [16]. These results suggest that a specific type of social prime can modulate automatic imitation; when individuals are personally primed to be prosocial, people find it harder to suppress their imitative tendencies.

Like imitation, accurate representation of another’s perspective is inherently intertwined with successful social interactions. Perspective taking has been shown to correlate with social competence [20] and successful communication requires both the ability to understand an interaction partner’s viewpoint and the ability to separate our own knowledge or beliefs from that point of view [21]. Perspective taking takes many forms, with visual perspective taking referring to situations where one must evaluate *what* someone else sees or *how* they see the environment [22]. Typically, individuals adopt an egocentric bias during social interactions, such that their own view is prioritised relative to others’ viewpoints [21, 23, 24, 25].

Unsurprisingly, such egocentrism can interfere with judgements about others’ perspectives [26, 27, 28, 29]. The Director’s Task [27, 28] requires participants to follow the instructions of another, the “Director”. In this task, a set of shelves, comprising sixteen slots (see Method section Figure 1), stand between the Director and a participant. The slots house a variety of familiar items (for example keys and cups), some of which are present in multiples of three and vary in size, and all of which were visible to the participant. However, a number of slots have a backing, such that any objects they contain are occluded from the Director’s view. The Director selects objects for the participant to remove from the shelves. On critical trials, the Director is not able to see the object that matches the description according to the participant’s view and it is on these trials that participants are required to deduce the item to which the Director is referring (e.g. select the second largest cup if the actual largest cup is not visible to the Director). The task indexes perspective taking by measuring the number of egocentric errors participants make on trials where there is a conflict between their perspective and the Director’s perspective. Even when it is made explicitly clear that the Director cannot not see the same objects that the participants can, egocentric errors are still made. That is, the participant responds as if the Director can see what they can. This suggests that while people may be capable of seeing things from another’s point of view, they do not always do so, with people often presuming another’s perspective is the same as their own [24,25]. As Gillespie and Richardson [25] put it; “although perspective taking is central to social life, people are not particularly good at it”. Identifying ways of improving its application should, therefore, enhance social interactions.

**Figure 1a:**
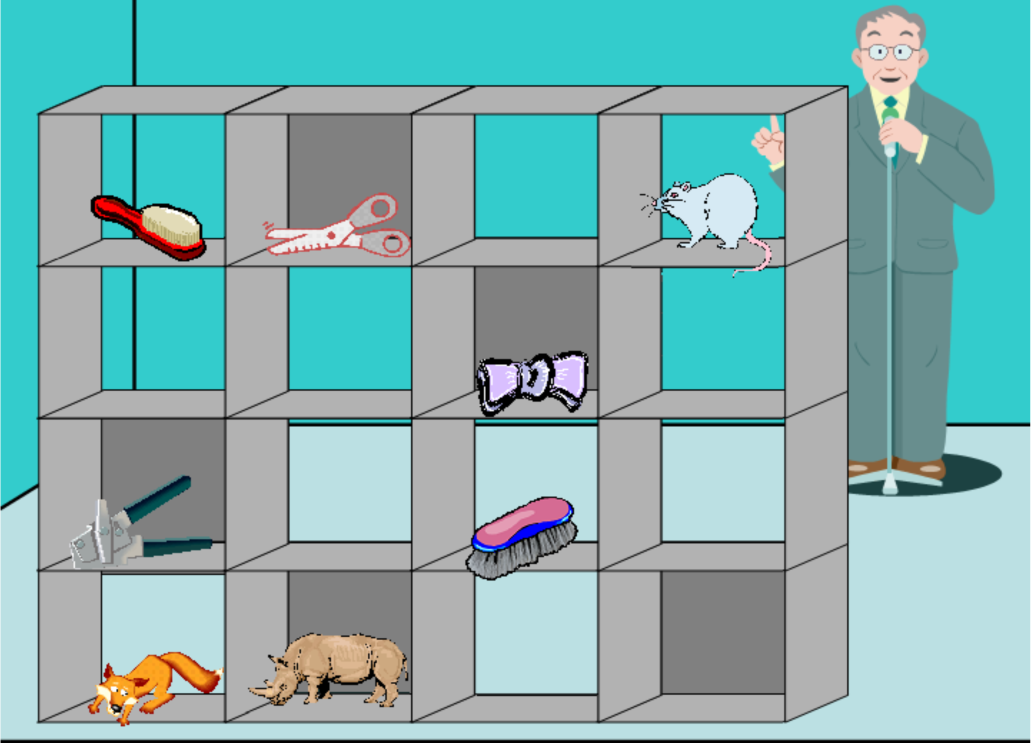
An example of a control trial (one item) in the Director’s task (“Move the mouse down”)

**Figure 1b:**
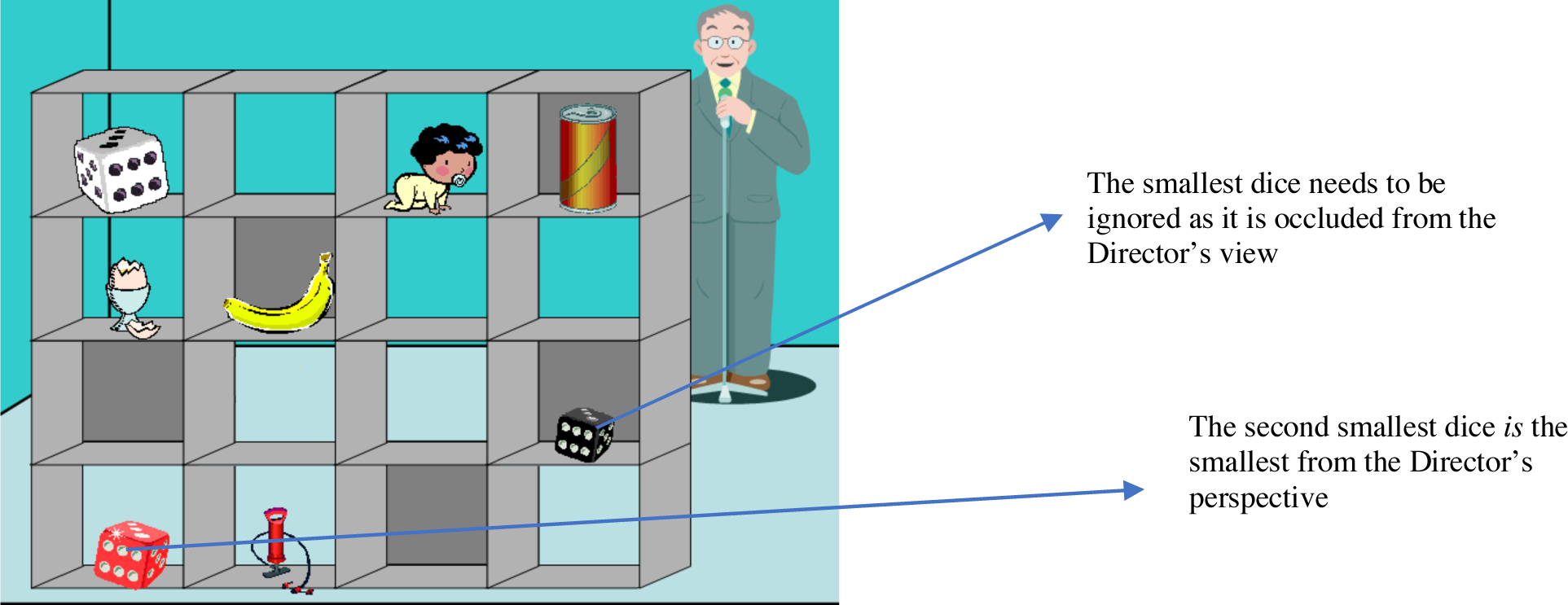
An example of an experimental trial in the Director’s task (“Move the small dice up”)

Although visual perspective taking has been studied extensively, how social context influences visual perspective taking and how visual perspective taking relates to other dimensions of social cognition, such as automatic imitation, have not been studied to date. Further, there is reason to suggest that automatic imitation and visual perspective taking may, in part, rely on a shared cognitive mechanism that distinguishes self from other. To succeed in automatic imitation tasks, a person must suppress the other’s action and promote their own. Conversely, for visual perspective taking, a person must suppress their own knowledge or belief and enhance the other’s perspective. Success at both tasks, then, requires a person to be able to quickly and flexibly distinguish between themselves and another. This is known as the ‘self-other distinction’ [see 30]. One study has directly addressed whether automatic imitation and visual perspective taking rely on a partially shared mechanism. Santiesteban and colleagues [31] found that training on a task that required a self-other distinction (imitation inhibition) transferred to a different self-other task; the Director’s Task. Even though automatic imitation and visual perspective taking may rely on a common mechanism, no research to date has shown that social context influences automatic imitation and visual perspective taking in a similar manner.

The current study, therefore, has three aims. First, drawing from studies exploring the effects of prosocial priming on automatic imitation, we will investigate the effects of prosocial priming on visual perspective taking. Does activating a goal to affiliate enhance one’s ability to readily adopt another’s visual perspective? Second, we will explore whether visual perspective taking and automatic imitation are correlated following prosocial priming. Does prosocial priming affect them in a similar manner? Third, we will perform a conceptual replication of previous studies, which showed an effect of first person, prosocial priming on automatic imitation [16, 17, 18]. Does activating a goal to affiliate increase automatic imitation in a subsequent RT task? Previous studies exploring this question have been conceptual replications of one another. While each used a different automatic imitation task, they all targeted and found the same main effect, indicating that the specific SRC task is not critical to the success of the prime. In addition, an effect was found irrespective of whether designs were within-subject [16] or between-subjects [17, 18] designs (see Method section, Table 3). Here then, a conceptual replication refers to studies using the same priming procedure to target the same effect while deviating on the precise automatic imitation task employed.

To test visual perspective taking abilities, we will use the Director’s Task [27, 28]. We will include both first person and third person prime conditions, to test whether self-relatedness influences prosocial priming of visual perspective taking in the same way as automatic imitation. Firstly, we predict that prosocially primed groups will achieve higher accuracy on the Director’s Task as compared to controls. Secondly, we predict that first person, prosocial priming will produce a positive correlation between visual perspective taking accuracy and larger congruency effects from the automatic imitation task. Finally, in line with previous findings, we expect that first person, prosocial priming will produce a larger congruency effect than both third person and control conditions. Together, these results will test the extent to which social context influences automatic imitation and visual perspective taking in a similar manner and therefore provide insight into the extent to which these core social abilities rely on a shared cognitive mechanism.

## Method

### Participants

Data from 153 individuals (111 female, mean age = 20.9, SD = 3.8, range 18-41) were collected in return for course credit; with 52 in the first person, prosocial (PS-1^st^) group, 52 in third person, prosocial (PS-3^rd^) and 49 controls. Ages ranged from 18 to 41 with average ages of 21.58 (SD 5.2) for PS-1^st^, 20.42 (SD 3.0) for PS-3^rd^ and 20.71 (SD 2.4) for the control group. Ethical approval was granted by the Research Ethics and Governance Committee of the School of Psychology at Bangor University. All participants gave their explicit informed consent and were free to withdraw from the study at any time.

### Sample Size & Power Calculation

No previous studies have explored the influence of prosocial priming on visual perspective taking, which means the expected effect size cannot be estimated from such data. Instead, the difference in congruency effects found by previous studies researching prosocial priming and automatic imitation (Table 3) was used to determine our sample size. These prior studies found medium to large effects (Cohen’s d of 0.53 - 0.75). However, evidence would suggest that published studies overestimate effect sizes [32, 33]. With this in mind, we powered our study to detect effect sizes at the lowest range of those found previously [34]. A sensitivity analysis in G*Power [35] using a one-tailed test, based on a mean difference between two independent groups (PS-1^st^ and control), with an alpha of .05 and 80% power to detect a medium effect size (Cohen’s d=0.5) or larger, returned a sample size of 50 participants per group. Therefore, we aimed to test 150 participants (50 per group) making our sample size much larger than previous studies (Table 3).

### Procedure and Stimuli

Prior to testing, participants were told they were taking part in a study investigating people’s accuracy rates and reaction time across three types of tasks. Testing was performed in one session, lasted approximately 45 minutes, and the order of tasks was held constant across participants. Upon arrival, participants were randomly assigned to a group; first person prosocial (PS-1^ST^), third person prosocial (PS-3^RD^) or control. The order of tasks was kept the same for all participants; first they completed a demographics information sheet and questionnaires, next they completed the priming task, immediately after priming they performed the perspective taking task and finally they completed the automatic imitation task. As our primary task of interest was the perspective taking task, we did not counterbalance the Director’s Task with the automatic imitation task as we did want any effects of imitation to confound any effects on perspective taking. Moreover, the Director’s task takes only around four minutes to complete (whereas the automatic imitation task takes over double that) meaning any effects of priming should survive the procedure.

#### 1. Demographics & Questionnaires

Prior to priming, each participant completed a brief demographics information sheet (age, gender, handedness and first language) together with three previously validated questionnaires; the Short Autism Spectrum Quotient (AQ-10 Adult) questionnaire [36], a self-esteem questionnaire [37] and the interpersonal reactivity index (IRI) [20]. Questionnaire data was collected for another study and is not discussed here. For completeness, the results are provided in Supplementary Table 1.

**Table 1:** Summary of accuracy (%) results from the Director’s Task. Mean accuracy (%) for control and experimental trials, together with overall accuracy, for each group are provided (sd in brackets)

|  | Trial Type |  | Overall Accuracy |
| --- | --- | --- | --- |
|  | Control | Experimental |  |
| PS-1<sup>st</sup> | 99.3 (1.7) | 97.3 (5.2) | 97.7 (2.8) |
| PS-3<sup>rd</sup> | 98.5 (3.6) | 97.2 (6.4) | 97.6 (3.8) |
| Control | 97.8 (4.5) | 95.7 (11.4) | 96.3 (5.2) |

#### 2. Pro-social Priming Stimuli

Prosocial priming was implemented through a scrambled sentences task [38] using sentence stimuli previously used to study automatic imitation [e.g. 16]. Three booklets, each containing 20 sentences, were used and each participant received only one booklet; either PS-1^ST^, PS-3^rd^ or the non-social control. Taking around 10-15 minutes, the task consisted of partially completed sentences with a list of words above them, with one word being irrelevant. Participants were instructed to select the correct words to write a grammatically correct sentence. PS-1^st^ and PS-3^rd^ sentences contained words such as together, collaborate, affectionate, share and help, which were designed to drive a prosocial attitude towards the self or the other respectively. All PS-1^st^ sentences started with ‘I’ whereas PS-3^rd^ used other people such that it was another person performing the prosocial act. For example, a completed first person, prosocial sentence might read “I always comfort my friends when they are upset” whereas the same sentence in the third person would read “David always comforts his friends when they are upset”. To produce a neutral attitude, control sentences were purely factual (e.g., “London is the capital of England”).

#### 3. Visual Perspective Taking

Following priming, the Director’s Task was administered. We used a computerized version of the Director’s Task [39], originally designed by Keysar and colleagues [27, 28]. The specific stimuli that we used were kindly shared with us by Dumontheil and colleagues [40]. Displayed on screen was a picture of a block of shelves (4−4 configuration) housing a number of recognisable objects, all of which were visible to the participant. Some shelves had a back on them such that anyone standing on the other side could not see the items in those slots (Figure 1). A person (the “Director”) was positioned on the other side of the shelves. The Director would issue an instruction (e.g. “Move the mouse down”) which the participant was required to follow by selecting the named object with the mouse and dragging it to the appropriate slot. Three practice trials were presented prior to the test beginning. Participants were explicitly made aware of the backing on some shelves and told that someone on the other side would not be able to see all of the items.

For the main task, there were 48 trials in total; 32 control trials (task involved only one object, which was visible to both participant and director, see Figure 1a), 8 non-conflict (NC) trials (more than one object of varying size, all visible to both participant and Director) and 8 conflict/experimental trials (more than one object of varying size, all visible to the participant but not all visible to the Director, see Figure 1b). To be correct on an experimental trial, the participant had to identify and move the object to which the Director was referring (see Figure 1b for an example). Trials were presented in blocks of three with participants only being given a short amount of time to respond before the next trial would automatically begin. The task was presented by ePrime version 2 and lasted for around four minutes.

#### 4. Automatic Imitation Task

Next, participants completed the automatic imitation task, based on the task designed by Brass and colleagues [11, 12]. Instructions were provided orally by the experimenter as well as in written form at the beginning of the task. At the start of each trial, participants were instructed to keep their index and middle fingers of their right hand pressed down on keys n and m respectively. Prior to each trial onset, the screen displayed a small fixation cross in the centre of the screen for 500ms. The image of a hand in a neutral position would then appear. Participants were instructed to raise their index finger when the number ‘1’ appeared on screen. When the number ‘2’ appeared, they were to raise their middle finger. Instructions were to respond as fast and as accurately as possible. To be correct on a trial, participants had to raise the finger that matched the number; index for ‘1’, middle for ‘2’. At the same time as the number appeared, the hand in the background would raise either its index or middle finger. For congruent trials, the stimulus hand would raise the same finger as the participant. For incongruent trials, the stimulus hand would raise a different finger to the participant (Figure 2).

**Figure 2:**
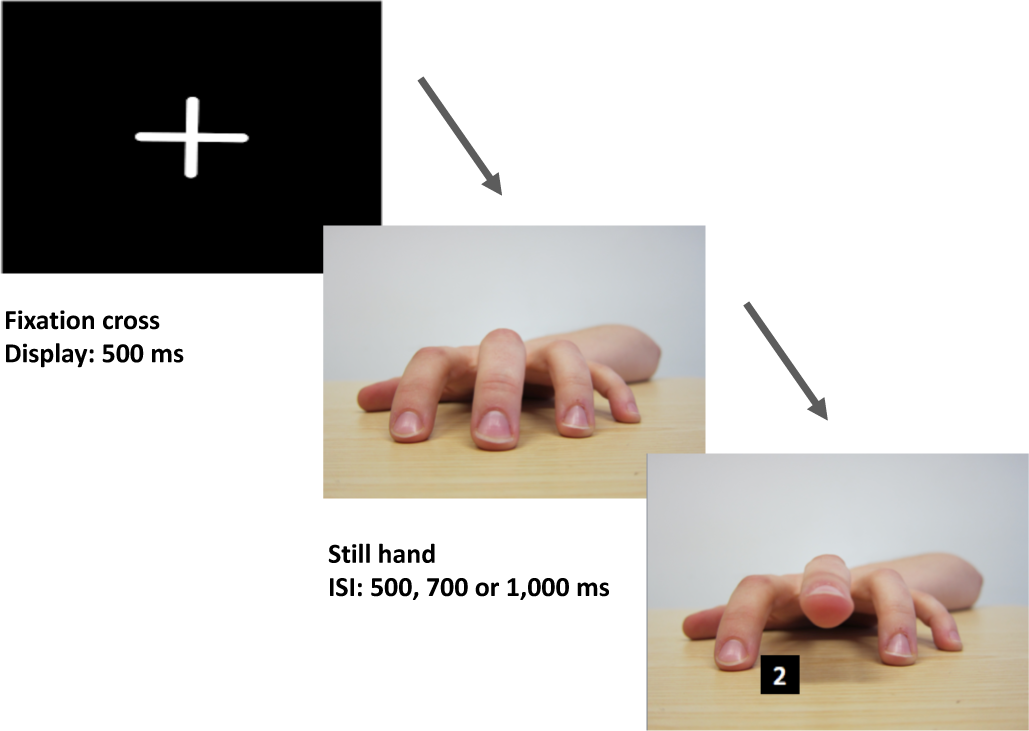
An example of a (congruent) trial in the automatic imitation task. The fixation cross is followed by an image of a still hand before the target stimulus (number) is displayed together with a lifted finger (irrelevant stimulus).

Data for 32 practice trials was collected prior to priming but not analysed. In the main task, there were 128 experimental trials in total, displayed in a random order, comprising 64 congruent trials (32 index and 32 middle) and 64 incongruent trials (32 index and 32 middle). Trials were presented in four blocks of 32 trials with an opportunity for a break being provided between each block. The task took around eight minutes to complete in total. In order to prevent participants from anticipating when the stimulus would appear, inter-stimulus intervals of 500, 700 and 1,000 milliseconds were randomly applied to the neutral hand before the next image appeared. The image of the hand and number would remain on screen until the participant lifted their finger or after 2,000ms, whichever came first, before returning to the fixation cross. The task was written in Matlab and presented using Psychophysics Toolbox.

Following completion of all tasks, participants were debriefed on the nature of the experiment. No participants reported guessing what the experiment was investigating and all were unaware that the scrambled sentences were trying to prime a prosocial attitude.

## Data analysis

### Visual Perspective Taking task – The Director’s task

In the version of the director’s task that we used, reaction time was not an instructive measure because the start point of the mouse curser was not fixed on every trial (i.e., participants could freely move the mouse during the instruction phase). As such, reaction time did not solely index the length of mental processing time; it also indexed the distance the hand had to travel to select the correct item. Reaction time data were not analysed. The accuracy of performance as a function of trial type and group was analysed. For each trial, participants could be correct, wrong or not answer (omit). Overall accuracy, based on correct responses for all 48 trials, was calculated for each participant. The mean accuracy and SD of each group was calculated. To control for outliers, participants with an average accuracy of less than three SD from their group’s mean were removed from their group. This resulted in seven participants being removed in total (PS-1^st^: 2; PS-3^rd^: 3; and Control: 2) and 146 being taken forward for analysis. For completeness, we also ran the analysis without removing outliers. Independent analysis of variance tests (ANOVAs) were used to explore differences in accuracy across the experimental groups.

### Automatic Imitation Task

In the automatic imitation task, reaction time was measured as the time taken from the appearance of the imperative cue (“1” or “2”) to when the finger was released. Trials were defined as accurate if the finger lifted matched the target number cue and incorrect if there was a mismatch between finger movement and target number cue. All incorrect responses were removed prior to analysis (<4% congruent trials and <10% of incongruent trials). Trials with a reaction time of less than 250ms or more than 2,000ms were also removed (<.1% of overall trials) as these were suggestive of expectancy errors and lapses in attention, respectively. Data for index and middle finger responses were collapsed. Accuracy and reaction time were calculated for each participant for each trial type; congruent and incongruent. Participants’ congruency effects were calculated by subtracting congruent reaction time from incongruent reaction time.

Outliers were considered in the context of both the individual (deviation from their own mean) and their group (deviation from the group mean). At participant level, trials falling outside of three SD either side of their mean reaction time were removed. Reaction time and accuracy for each participant was recalculated and taken forward into the group calculations. Group reaction time and accuracy means were then calculated and participants falling outside of three SD of their group’s mean (for either reaction time or accuracy) were removed from further analysis. This resulted in six participants (PS-1^ST^: 1; PS-3^RD^: 1; and control: 4) being removed from further analysis and 147 being taken forward. ANOVA was used to test for differences in accuracy, reaction times and congruency effects across the experimental groups. To ensure that the removal of outliers did not affect the outcome of our results, analyses were repeated on the complete dataset.

## Results

### Visual Perspective Taking Task

Accuracy for all trial types across all groups are reported in Table 1. Performance on the task was high across all groups, with average accuracy exceeding 90% for experimental trials (Figure 3). Errors on experimental trials were rare and trials that were omitted (left unanswered) were more common (Figure 4). This would suggest that, of the trials completed, there was a ceiling effect present in performance (117 participants scored 100%, 26 scored 87.5% and the remaining 10 scored 75% or less). Accuracy for control and experimental trials (conflict between participant’s and Director’s perspective) were compared between groups. Using group as the between subject’s factor, two one-way ANOVAs on trial type revealed no significant differences between groups for accuracy on control F(2,143)= 2.31, p=.103, np^2^=.031 or experimental (F(2,143)= 0.53, p=.587, np^2^=.007) trials.

**Figure 3:**
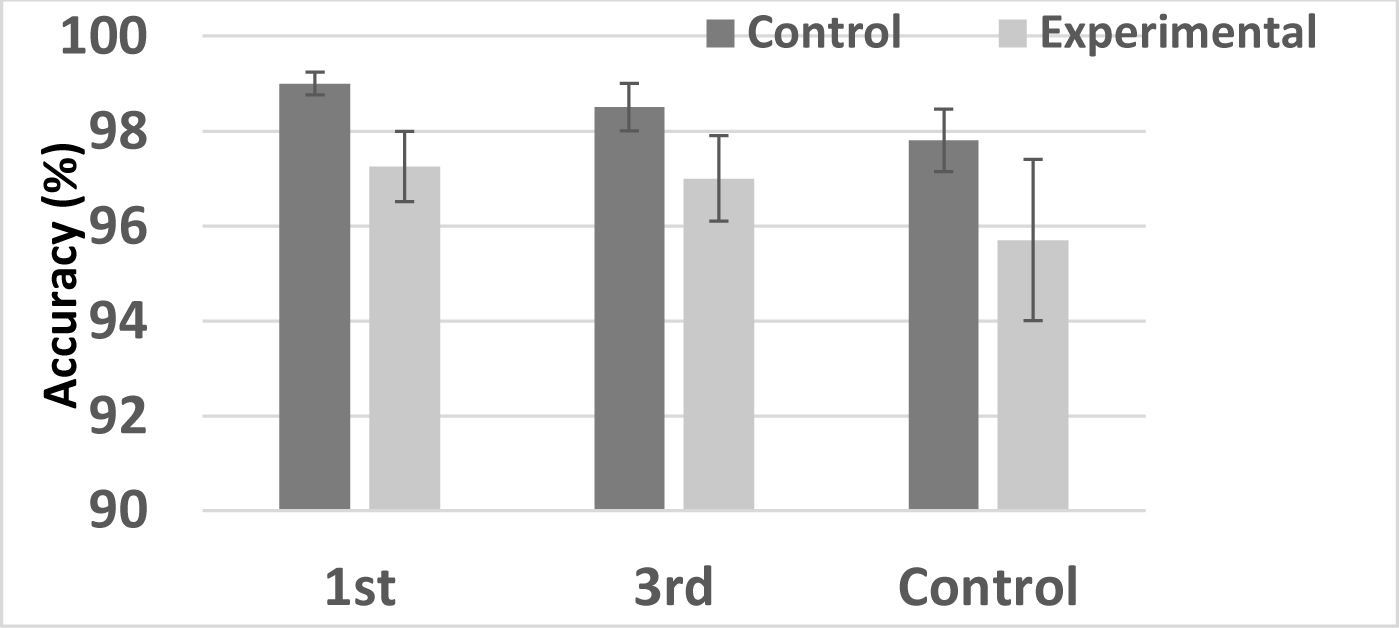
Accuracy (%) for control and experimental trials on the Director’s Task for each group. Bars represent SEM

**Figure 4:**
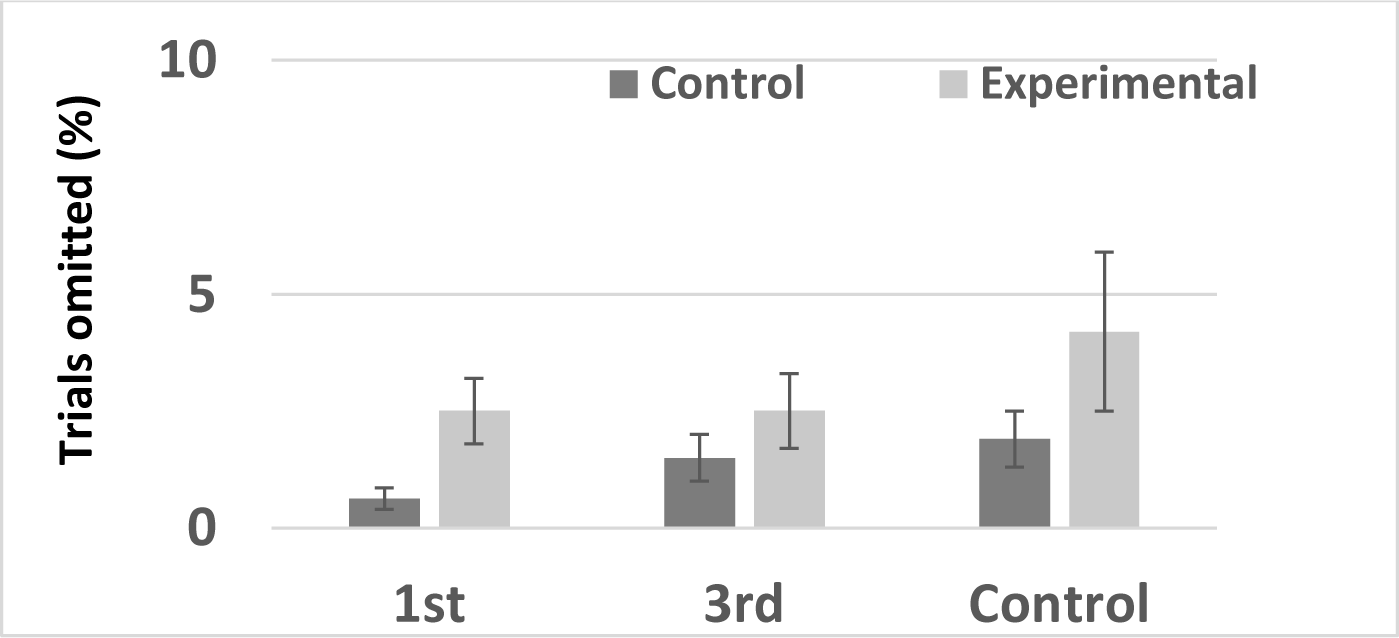
Omissions (%) for control and experimental trials on the Director’s Task for each group. Bars represent SEM

Given the overall high accuracy across all groups, which is indicative of a ceiling effect, further analyses of the relationship between visual perspective taking automatic imitation were not performed as they would not be instructive.

The Director’s task was used because many studies report substantial error rates when using it and, as such, a ceiling effect was not expected. Near perfect scores across all experimental groups in this study prompted a (non-exhaustive) review of studies using the same task with adult participants (Supplementary Table 2). The search revealed that the task returns a variety of results ranging from 54-88% accuracy. Worth noting is the fact that the task only includes eight experimental trials, thus this range translates to one to four errors. For instance, accuracy of 87.5% (7/8) would be achieved if only one mistake was made.

**Table 2:**
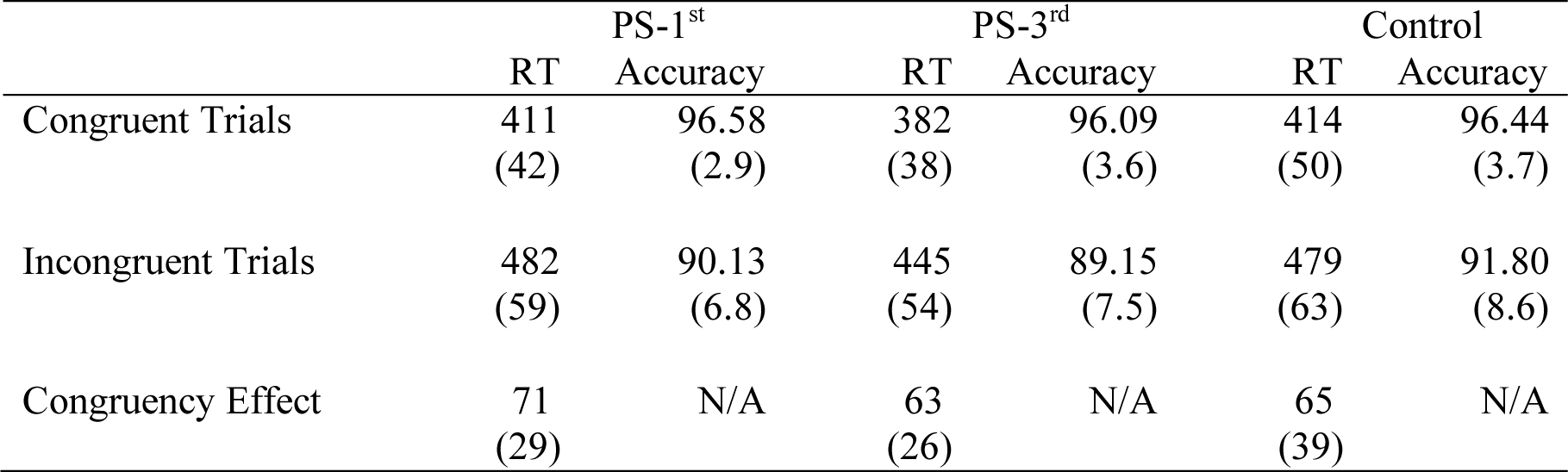
Summary of results from the automatic imitation task with reaction times (ms) and accuracy rates (%) for each trial type and the congruency effect (incongruent RT – incongruent RT) for each group (sd in brackets)

### Automatic Imitation task

Mean reaction times for congruent and incongruent trials, as well as the congruency effect (CE) are reported in Table 2. As can be seen, participants were faster and more accurate on congruent trials (Figures 5 and 6). A repeated-measures ANOVA was performed with congruency (trial type: congruent and incongruent) as the within-subjects factor and group (PS-1^st^, PS-3^rd^ and control) as the between-subjects factor. There was a significant main effect of congruency F(1,144)=647.759, p<.001, ηp^2^=.818, with congruent trials being significantly faster than incongruent trials. There was also a significant effect of group F(2,144)=7.882, p=.001, ηp^2^=.099. Mean RTs for each experimental group (collapsed across congruent and incongruent trials) were calculated and compared using t-tests. These analyses showed that the PS-3^rd^ group was significantly faster than both the PS-1^st^ t(100)=3.65, p<.001 and control t(94)=3.32, p=.001 groups (see Figure 3). There was no mean RT difference between the PS-1^st^ and Control group t(94)=.004, p=.997. Crucially, there was no interaction between congruency and group F(2,144)=0.943, p=.392, ηp^2^=.013 indicating there was no differential effect of priming on congruency between groups.

**Figure 5:**
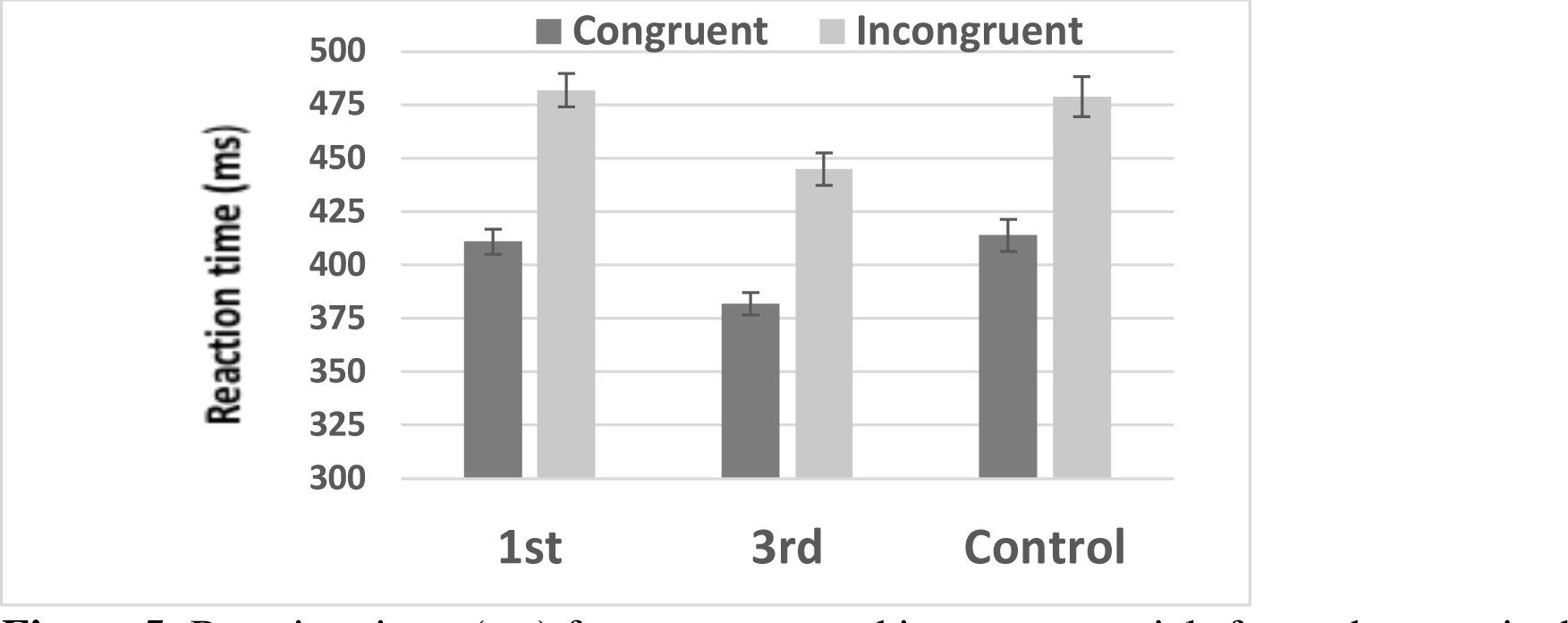
Reaction times (ms) for congruent and incongruent trials for each group in the Automatic Imitation Task. Bars represent SEM

**Figure 6:**
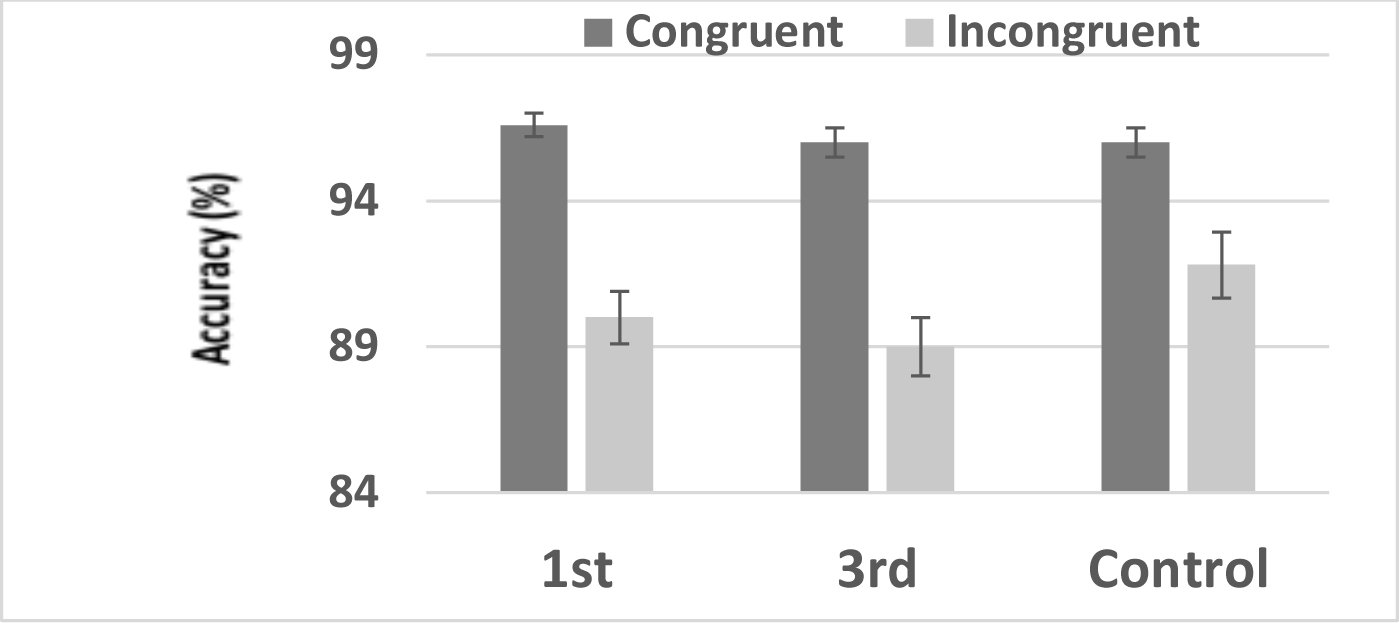
Accuracy rates (%) for congruent and incongruent trials for each group in the Automatic Imitation Task. Bars represent SEM

As prior studies analysed the congruency effect [16, 17, 18] we carried out an independent one-way ANOVA on congruency effect as a function of group (Figure 7). There was no significant difference between the groups’ congruency effects F(2,144)=0.96, p=.387, ηp^2^=.013. To ensure that the removal of outliers had not changed the results, we ran the same test with all participants (except for one who did not complete the task) included. The result was the same F(2,149)=1.24, p=.291, ηp^2^=.016. In addition, we wanted to ensure that English language proficiency did not impact priming effects. When removing non-native English speakers (N=29), there was still no effect of priming on imitation F(2,121)=1.2, p=.304.

**Figure 7:**
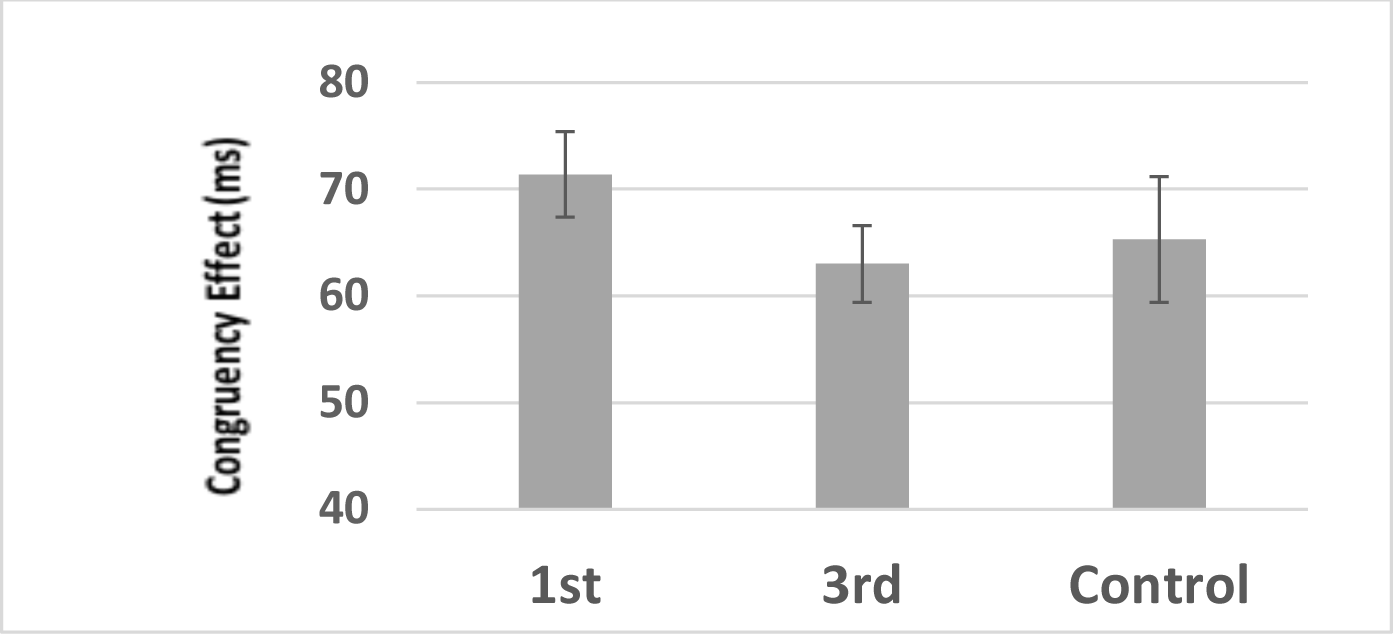
Congruency Effects (CE) – incongruent RT less congruent RT – for each group for the Automatic Imitation Task. Bars represent SEM

To provide quantitative evidence for the null hypothesis, a Bayesian analysis was performed [41] in JASP using the independent t-test function [42]. The returned Bayes factor BF^01^ provides an estimate of how likely the null hypothesis (0) is compared to the experimental hypothesis (1), given the data. A Bayes factor of 3.3 was returned. This suggests that the null hypothesis was three times more likely than the experimental hypothesis [43].

### Meta-Analysis of automatic imitation results: PS-1^st^ vs Control groups

To put our automatic imitation result in context, we performed a meta-analysis. The three previous studies using first person, prosocial priming (scrambled sentences) to investigate the effects on automatic imitation were included in the meta-analysis, along with the current study (Table 3). While these studies covered both within- [16] and between- [17, 18] subject designs and employed slightly different methods for testing automatic imitation, they shared sufficient similarity to be directly compared. All four studies used scrambled sentences to prime prosociality and measured imitation via an SRC index of automatic imitation. Therefore, while these studies are not direct replications of each other, they have substantial methodological similarity and all target the same primary effect, such that we consider them conceptual replications of each other. We meta-analysed the difference in congruency effect for first person priming compared to control. We were able to obtain raw data from one study [16]. In the absence of raw data for all studies, we used the values available from the published studies to compute standard deviations, standard errors and effect sizes. Cohen’s d [44] was calculated as the mean group difference divided by the pooled standard deviation.

**Table 3:** Summary of studies included in the meta-analysis. Mean congruency effects (CE) for PS-1^st^ and control groups (sd in brackets) are used to calculate the standardised effect size (Cohen’s d). (* PS1st (51) and Control (45) were different sample sizes). Spatial stimuli introduce both a spatial and imitative component to the design. Orthogonal stimuli rotate the stimuli to reduce (but not remove) the spatial component.

| Study | Design | Stimuli |  | Sample/<br>Group<br>size | PS-1<sup>st</sup>(2)<br>CE | Control (1)<br>CE | Effect Size (d)<br>(2-1)/pooled sd |
| --- | --- | --- | --- | --- | --- | --- | --- |
| Wang & Hamilton (2013) | Within | Fingers | Spatial | 16 | 28 (16) | 16 (16) | 0.75 |
| Cook & Bird (2011) | Between | Fingers | Orthogonal | 28 | 71 (63) | 38 (37) | 0.66 |
| Leighton et al (2010) | Between | Hands | Spatial | 12 | 38 (31) | 26 (14) | 0.53 |
| Current study | Between | Fingers | Spatial | 45-51* | 71 (39) | 65 (29) | 0.18 |

The meta-analysis was performed using Exploratory Software for Confidence Intervals [45] ESCI calculates a weighted contribution for each study based on sample size and variance, with larger sample sizes and smaller variance receiving the highest weighting. Based on Cumming’s recommendations [45], we used a random effects model to estimate the likely population effect size in original units (ms), as well as standardized units. 95% CIs are reported as a measure of precision for these population estimates. The results from these two calculations are reported here using forest plots (Figures 8a and 8b).

**Figure 8a:**
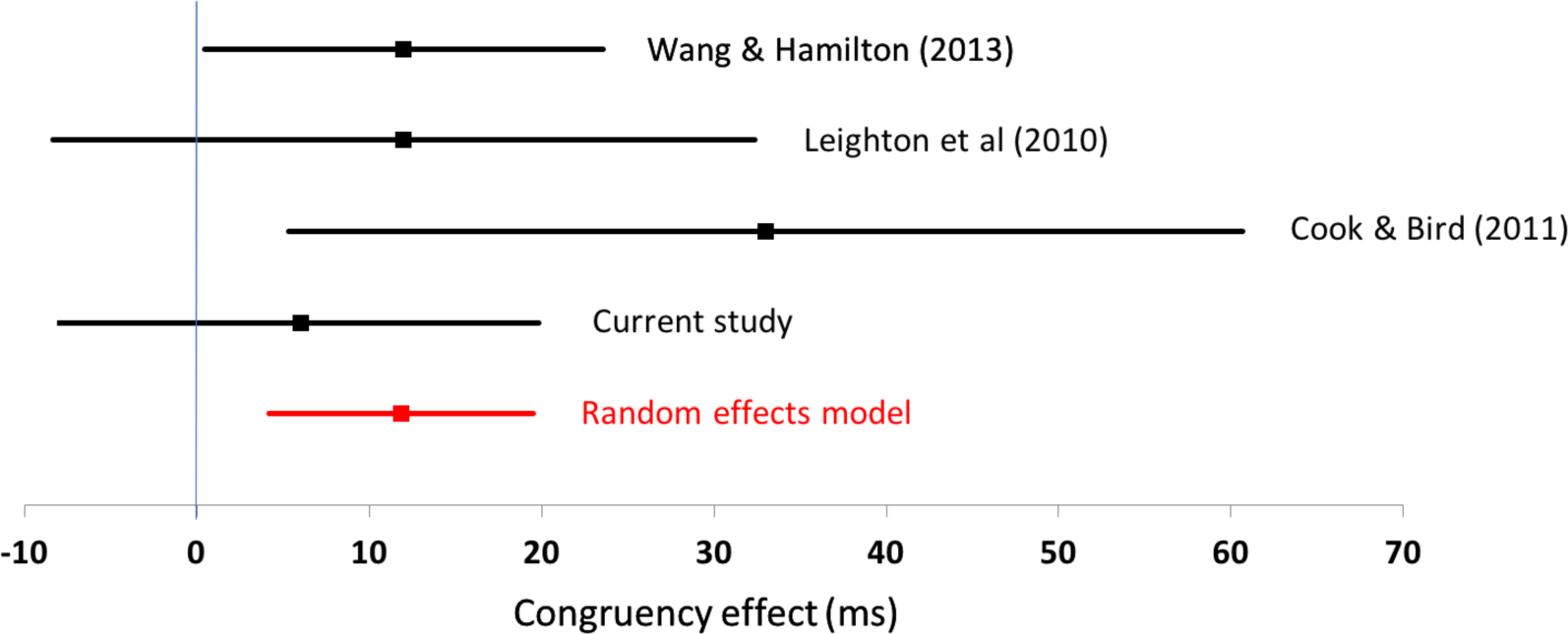
Forest plot of meta-analysis for Original units (ms). Lines represent 95% confidence intervals. The random effects model indicates the likely population effect.

**Figure 8b:**
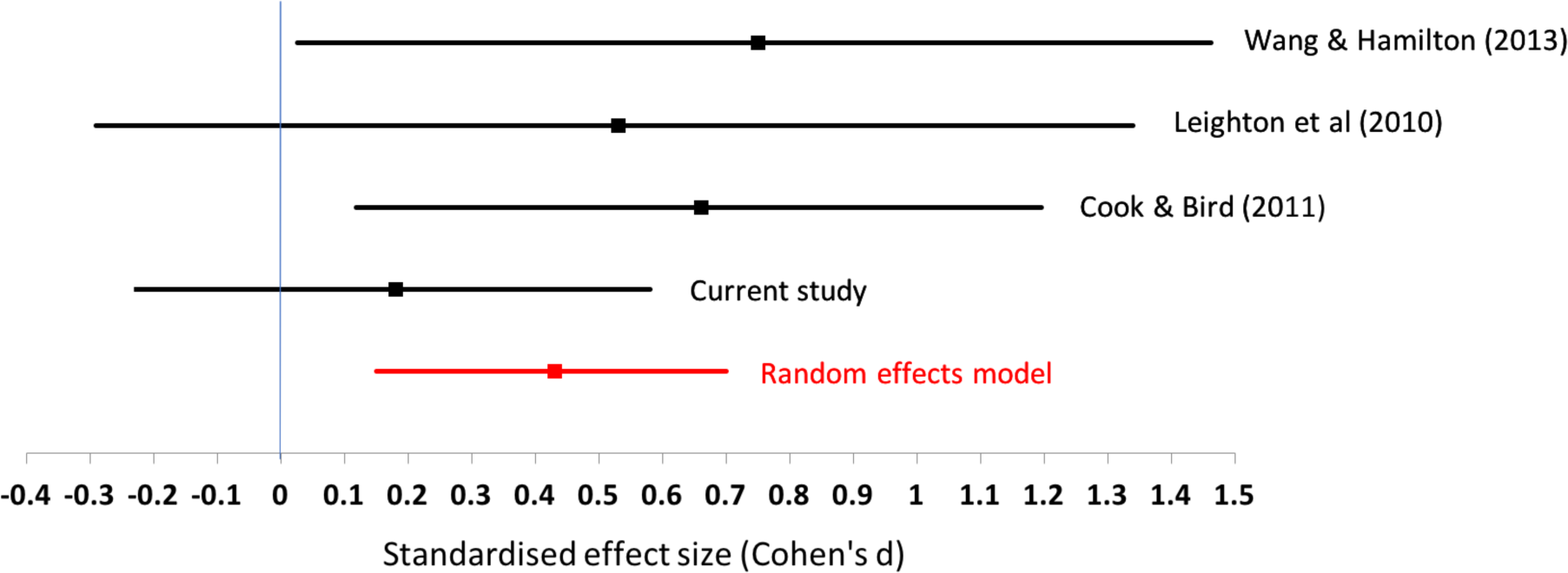
Forest plot of meta-analysis for Standardised units (Cohen’s d). Lines represent 95% confidence intervals. The random effects model indicates the likely population effect is somewhere between a very small and medium effect.

The estimated difference in priming between first person and control is 11ms [95% CI 4, 19] (Figure 8a). As can be seen from Figure 8a, two of the four studies in the MA have confidence intervals (CI) that cross over the zero line and the effect sizes range from 4 to 19ms. The standardized effect size is d=0.43 [0.15, 0.7] (figure 8b), and varies across the four studies, with interval estimates touching or crossing zero in three of the studies. These results suggest that the effect is imprecise and it is possible that the true effect size may be close to zero. Prior to running this study, the cumulative effect size based on three prior studies was d=0.64. Adding the current study, which has a much larger sample size than all prior studies, reduces the cumulative effect size by a third to d=0.43 (Figure 8b).

### Open data

To aid future meta-analyses and power estimates, data from the current experiment are available online for all dependent measures (osf.io/bseky).

## Discussion

Due to a ceiling effect using the Director’s Task, we were unable to investigate the effects of prosocial priming on visual perspective taking. A comparison of other studies using the Director’s Task shows wide variability in accuracy scores. Accordingly, we suggest that the reliability of the measure may be low and future research should test this formally. In addition, and contrary to previous studies and our expectations, we found no effect of prosocial priming on automatic imitation. To better understand this unexpected result, we performed a meta-analysis of the effects of prosocial priming on automatic imitation. The result indicates that if a relationship does exist between prosocial priming and automatic imitation, it is likely smaller and more variable than the results of any one previous study would suggest. Therefore, we offer caution when using the Director’s Task as a measure of perspective taking and reduce the strength of evidence in favour of social priming modulating automatic imitation. More generally, the current study demonstrates the utility of replicating and meta-analysing main effects in an effort to build a more cumulative science of social cognition.

### Prosocial priming and Visual Perspective Taking

We found an unexpected ceiling effect in the Director’s Task and, therefore, could not perform our primary analyses of interest. We reviewed published studies that have administered the Director’s Task to adults (over 18) and reported their accuracy rates (Supplementary Table 2). This brief review found that the task returns a range of results (54-88%). As a consequence, and in addition to concerns over the validity of the Director’s Task as a measure of visual perspective taking [46, 47, 48, 49], these findings suggest that the Director’s Task could have low reliability, such that task performance appears to vary quite substantially from study to study. As such, we recommend that future studies should formally evaluate the reliability of the measure before using it further.

We also note other features of the Director’s Task that are worth further consideration in future research. Not all studies using the Director’s Task specifically state the number of trials analysed, so it is possible that accuracy scores vary across studies because of methodological differences in the way the task was administered. Further, when interpreting accuracy scores, it is important to note that there are only eight experimental trials; a factor we did not fully consider when designing the study. Scores of 75% and 87.5% may seem substantially different, but in this task, the difference is only one error. This does not bode well for studies such as ours, which aim to improve perspective taking scores through experimental manipulations or training (in this case, through prosocial priming). There simply is not enough “room” to measure any true increase in the skill with adult participants. It could be argued that more trials are needed in the experimental condition, however, given the accuracy rates returned in our data, participants seem to reach ceiling quickly, rendering the data from those extra trials superfluous. Therefore, we offer caution to those interested in visual perspective taking in using the Director’s Task, especially if the research question relies on score variability or score manipulation.

### Prosocial priming and Automatic Imitation

Previous studies have shown that PS-1^st^ priming leads to increased congruency effects on automatic imitation tasks [16, 17, 18]. Although the current study had the power to detect effects smaller than those previously observed, we did not observe an effect. While we did find a small reaction time difference (6ms) between the PS-1^st^ priming and control groups in the same direction as previous studies, the difference was not distinguishable from zero. Further, a Bayesian analysis provided three times more support for the null over the experimental hypothesis.

Of the four studies included in the meta-analysis, one has a 95% confidence interval that touches the zero line and two actually cross the line (Figure 8b). This is suggestive of an imprecise estimate of a population effect size, which could be small in size (close to zero) and paints a different picture to the way in which effects were interpreted by each individual study. Overall, the pattern of results is in keeping with suggestions in the literature that published effects are commonly over-estimated [32, 33] and underscores the value of meta-analytic thinking when aiming to synthesise prior findings [45, 50]. It is more than likely that the actual effect of prosocial priming on automatic imitation is smaller than previously reported as the meta-analysis suggests a population effect size of d=0.43. The meta-analysis also illustrates the variability of findings to date, with confidence intervals for the standardised effect size ranging from 0.15 to 0.70. In addition, viewing our null result (d=0.18) within the context of the meta-analysis (d=0.43) suggests that the effect of first person, prosocial priming on automatic imitation is indeed prone to variation.

### Limitations and future directions

One potential limitation of the current study is the imitation task used has a spatial compatibility component, which might introduce ‘noise’ to the data that could interact with the imitative tendencies of the participants [2, 51, 52]. Although possible, it is unlikely to have been the reason behind our null results. Prior studies used the same task and were able to show effects of the same social priming technique on congruency effects [16]. Therefore, while we do not think it can account for the current null results, separating imitative tendencies from spatial compatibility would be a useful future direction for research investigating automatic imitation more generally [51, 53, 54].

One further limitation concerns the sequencing of tasks. To avoid any influence of the imitation task on the Director’s Task, we used a fixed order across participants. It is therefore possible that, by administering the Director’s Task prior to the automatic imitation task, we unwittingly introduced another prosocial prime that interfered with the effects of the intended prosocial prime. That is to say, taking someone else’s perspective may itself serve as a prosocial prime that increases the tendency to imitate. However, if the prosocial prime and the visual perspective taking task both activated a goal to affiliate, we might still expect to observe overall greater imitative tendencies in the first person, prosocial group; the effects on behaviour from both primes might be expected to be additive. This possibility is not supported by the current data due to the fact that the control group returned the same congruency effects as the prosocially primed group.

Conversely, if participants did have a goal to act prosocially, the completion of the Director’s task could have satisfied this goal and, in essence, ‘switched off’ the prime. Again, this explanation could account for the lack of group differences in the automatic imitation task as all groups could have been returned to baseline. If this were the case, it would still not encourage thinking of priming as a robust method for increasing prosocial behaviour; as soon as one completes the goal, they return to a neutral position. Alternatively, the visual perspective taking task could have diluted, or even overwritten, any effects the prosocial priming task may have generated, which would account for the lack of group differences. However, with only eight trials among 48 actually requiring the participant to take someone else’ perspective, the visual perspective taking task would need to exhibit strong effects to remove those created by the prosocial priming task administered just five minutes previously. Ultimately, a future study is required to determine whether the Director’s Task can function as a prosocial prime that modulates imitative tendencies.

In summary, the order effect created three possibilities that could in theory account for this study failing to find the same effect on automatic imitation following prosocial priming as that found by other studies. Either 1) the goal from priming was satisfied by completing the Director’s task, 2) the Director’s task exerts effects strong enough to return all groups to baseline (or equally primed) or 3) the effects of prosocial priming are too weak to survive an intervening task. While no firm conclusions can be drawn at this moment, when considering these possibilities and the highly variable effect highlighted by the meta-analysis, it is prudent to say that the influence of prosocial priming on automatic imitation is unlikely to be robust.

### Conclusion

Due to an unforeseen ceiling effect in the Director’s Task, we could not evaluate whether prosocial priming modulates visual perspective taking and this question remains open for future studies to address. Instead, we suggest that when investigating visual perspective taking using the Director’s Task, the possibility that the task has low reliability with adult populations should be given due consideration and formally tested. The current study also questions the robustness of prosocial priming effects on automatic imitation. Indeed, meta-analysing all studies to date suggests that the effects of prosocial primes on imitation are variable and could be small. Finally, by reporting null results we hope to avoid the file drawer problem and inherent bias in the published literature [55, 56]. Also, by meta-analysing results as studies emerge [45, 50] and by making raw data freely available [57], we hope to move towards a more cumulative science of social cognition that future studies can build upon.

## Acknowledgements

This research was performed as part of an all-Wales ESRC Doctoral Training Centre 1+3 PhD studentship.

## Compliance with Ethical Standards

Conflict of Interest: All authors declare that they have no conflicts of interest. Ethical approval: The data reported here were obtained from human participants under approval from the Research Ethics and Governance Committee of the School of Psychology at Bangor University.

Informed consent: Informed consent was obtained from all individual participants included in the study.

